# [^18^F]-Labeled PARP-1 PET Imaging of PSMA Targeted Alpha Particle Radiotherapy Response

**DOI:** 10.1101/2022.03.16.484613

**Authors:** Hanwen Zhang, Diane Abou, Alexandria Villmer, Nadia Benabdallah, Wen Jiang, David Ulmert, Sean Carlin, Buck E. Rogers, Norman F. Turtle, Michael R. McDevitt, Brian Baumann, Brian W. Simons, Dong Zhou, Daniel L. J. Thorek

**Author notes:** co-corresponding authors Radiological Sciences, Washington University School of Medicine, 510 S. Kingshighway Blvd. Campus Box 8225, St. Louis, Missouri, 63110, |.

## Abstract

**Motivation:** The growing interest and clinical translation of alpha particle (*α*) therapies brings with it new challenges to assess target cell engagement and to monitor therapeutic effect. Noninvasive imaging has great potential to guide *α*-treatment and to harness the potential of these agents in the complex environment of disseminated disease. Poly(ADP) ribose polymerase 1 (PARP-1) is among the most abundantly expressed DNA repair enzymes with key roles in multiple repair pathways - such as those induced by irradiation.

**Materials and Methods:** Here, we used a third-generation PARP1-specific radiotracer, [^18^F]-PARPZ, to delineate castrate resistant prostate cancer xenografts. Following treatment with the clinically applied [^225^Ac]-PSMA-617, positron emission tomography was performed and analyzed, along with and correlative autoradiography and histology acquired.

**Results:** [^18^F]-PARPZ was able to distinguish treated from control (saline) xenografts by uptake. Kinetic analysis of tracer accumulation also suggests that the localization of the agent to sites of increased PARP-1 expression is a consequence of DNA damage response.

**Conclusions:** Together, these data support expanded investigation of [^18^F]-PARPZ to facilitate clinical translation in the *α*-therapy space.

## INTRODUCTION

Radiation is a mainstay of cancer therapy, most often generated by linear accelerator systems with image guidance. Indeed, approximately 50% of all cancer patients will receive radiation of some form during their treatment. In the metastatic setting, radiation delivery becomes much more complex as planning for treatment at multiple foci must also ensure that neighboring healthy tissues do not receive ablative doses. Thus, radiation to treat disseminated disease is less frequently applied. The use of targeted radiation delivery from beta or alpha particles that are administered systemically and localize to deposits of disease overcomes several of these issues.

Alpha particles (*α*) are of particular interest as these high-energy, high linear energy transfer helium nuclei deliver tumoricidal doses with emissions that are highly localized^1^. Recently, Radium-223 dichloride, the first approved drug in this class, was marketed for the management of bone metastases in castrate resistant prostate cancer patients^2^. This follows a successful Phase III clinical trial which demonstrated an overall survival advantage for patients receiving the bone-targeted agent^3^. The approval of Radium-223 and the initial clinical investigation of other targeted alpha particle (*α*)-therapy vehicles^4–7^, has led to an increased interest in the utilization of these potent emitters.

The double strand DNA breaks and severe genotoxic damage incurred by *α*-traversal of the nucleus are difficult to repair. In controlled *in vitro* study, a single *α* can kill a target cell^8^. However, the short path length of these emission means that many cells are spared any exposure; as evinced in part by the fact that these are not curative treatments^3,9^. Furthermore, we have shown previously that genomic factors such as DNA damage repair (DDR) status may influence sensitivity to *α*-therapy^10^. We and others posit that detection of DDR, ideally through non-invasive methods, can be used to personalize treatment approaches^11–14^.

The multipathway mechanisms of eukaryotic DDR are intricate, orchestrated and tightly controlled processes. Summarily, these involve sensing of damage, signal transduction and recruitment of enzymatic complexes to remodel and repair DNA. Poly (ADP-ribose) polymerase 1 (PARP-1) is among the most well studied DDR proteins^15–17^ and has critical roles in multiple single and double strand repair processes^18^. As such, inhibition of catalytic activity of this enzyme, and in particular the PARP-1 isoform, has been exploited to induce synthetic lethality in combination with loss of BRCA1 or BRCA2 function. The capacity to assess the PARP-1 status in a target cancer cell is emerging as a means to select patients for and to monitor PARP-1 inhibitor therapy^19,20^.

[^18^F]PARPZ, also known as [^18^F]WC-DZ-F, was developed as an analogue of [^18^F]FluorThanatrace (FTT) and [^125^I] KX1 which are well-established PARP-1 ligands for measuring PARP-1 expression. Our previous work has shown that this third-generation radiolabeled PARP-1 inhibitor has highly specific targeted uptake and favorable pharmacokinetic properties for use as a non-invasive imaging agent and targeted therapy^21^. In this study, we have evaluated the application of [^18^F]PARPZ in a model of BRCA1 null castrate resistant prostate cancer (22Rv1) to detect DDR to in vivo *α*-irradiation using [^225^Ac]-PSMA-617 (Figure 1). This chelate-conjugated peptide targets prostate specific membrane antigen (PSMA, also known as Glutamate carboxypeptidase II [GCPII]), a cell surface receptor of considerable clinical and research interest in prostate cancer^4,22,23^.

**Figure 1:**
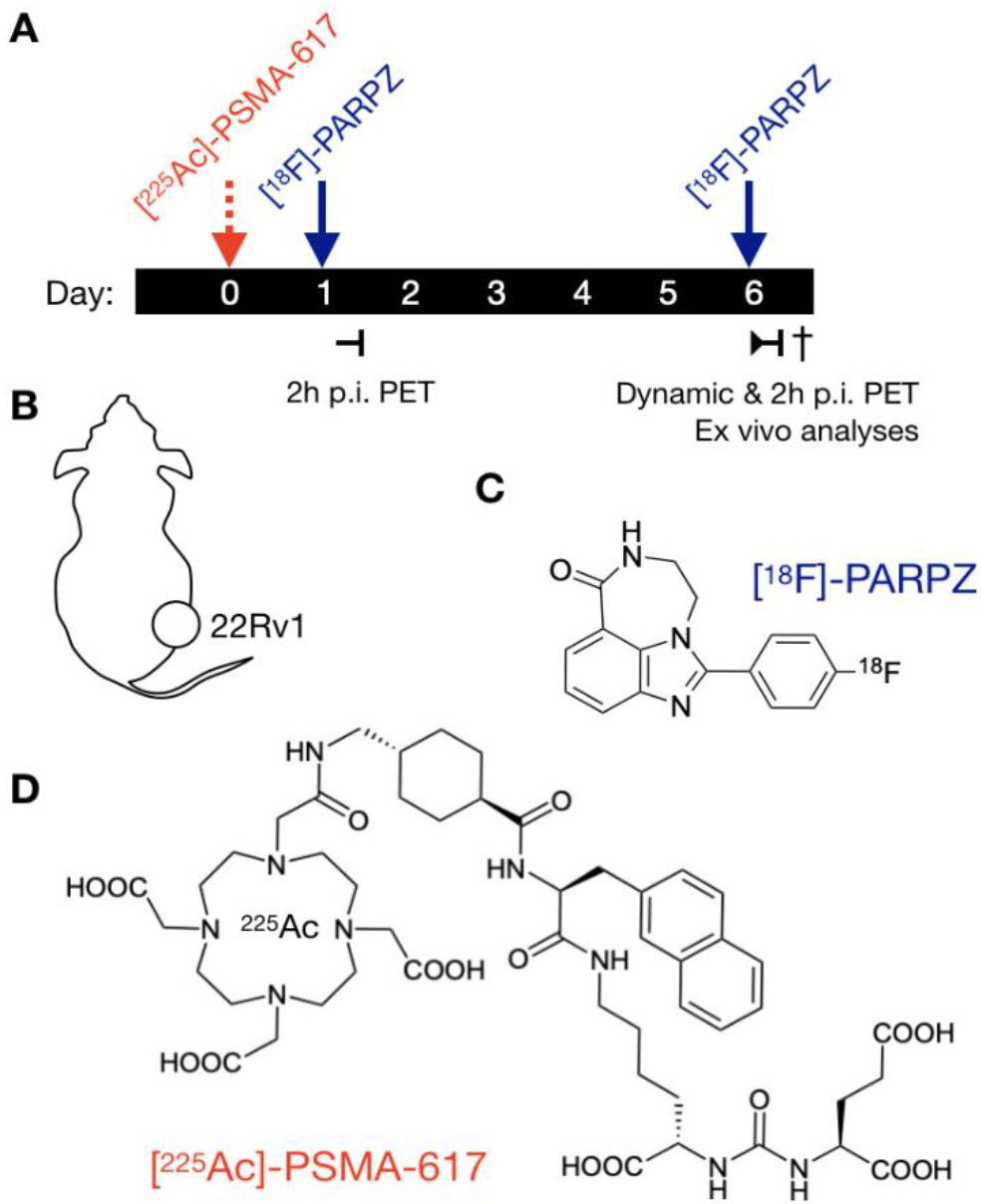
Schematic of Study Design. A) PSMA-expressing 22Rv1-Luc subcutaneous xenograft were implanted into NCI athymic Nu/Nu male. Upon reaching approximately 500 mm^3^, either control saline or the potent targeted [^225^Ac]-PSMA-617 was administered. Static [^18^F]PARPZ-PET was performed at 1 and 6 d post-therapy (n=8), and dynamic imaging for group matched animals (n=3). Chemical structures of A) the PARP-1 specific [^18^F]-PAPRZ radiotracer and B) the prostate cancer targeting [^225^Ac]-PSMA-617.

## MATERIALS AND METHODS

### Synthesis and Radiosynthesis of [^18^F]PARPZ

[^18^F]PARPZ was synthesized and prepared for injection as previously described^21^, with improvements. [^18^F]-PARPZ was synthesized by two steps according to previously reported procedure. Synthesis of 4-[^18^F]fluorobenzaldehyde ([^18^F]**2**): Into a 10 mL Pyrex tube containing 4-formyl-N,N,N-trimethylbenzenaminium triflate precursor (3.2 mg, 10.2 µmol) and K_2_CO_3_/K_222_ (2.8 mg, 3.1 µmol) was added [^18^F]tosyl fluoride (1.32GBq/35.7 mCi) in acetonitrile (0.5 mL). The reaction mixture was shaken and then heated in an oil bath at 108 °C for 7 min. Synthesis of [^18^F]-PARPZ: At room temperature, the above reaction mixture passed through a Waters MCX cartridge (100 mg in 1 mL cartridge, pretreated with 3 mL acetonitrile), following by acetonitrile (0.5 mL) for rinsing, and all were eluted into a 10 mL Pyrex tube containing precursor **1** (2.6 mg, 14.7 µmol)) and 10% Pd/C (6 mg) in methanol (0.5 mL). The tube was flushed with argon, and then capped firmly and heated at 120 °C in a heating block. After 20 min, the reaction was completed according to HPLC. The reaction mixture passed through a pad of Celite (pre-treated with acetonitrile), and the reaction tube and Celite were rinsed with acetonitrile (2 mL). All the eluted solution was collected in a tube, and solvents were removed under a flow of argon at 108 °C. The residue was dissolved in HPLC mobile phase/water 1:1 (4 mL), and injected onto HPLC via a nylon filter. After HPLC purification (Agilent SB-C18 250×9.4 mm, 17% acetonitrile/83% water/0.1% TFA, 4 mL/min, 250 nm), [^18^F]-PARPZ (0.44 GBq/12 mCi) was collected at 11 min and diluted in water (40 mL). [^18^F]-PARPZ was extracted from the solution by a standard solid phase extract procedure using a Waters HLB light and was formulated in 10% ethanol/saline.

### Radiolabeling of [^225^Ac]-PSMA-617

Actinium-225 was supplied as a dry nitrate (Oak Ridge National Laboratory, Department of Energy). [^225^Ac]-PSMA-617 was prepared by dissolving 10 μg peptide in 97.5 μL 0.25 M HEPES buffer (pH 8.6), and adding 185 KBq [^225^Ac](NO_3_)_3_ solution (2.5 μL) followed by a 15 min incubation at 97 °C. After incubation, the reaction solution was analyzed with TLC (C18; AcCN/10 mM EDTA buffer (1/9) as mobile), and the labeling yield was greater than 95%. After purification with Strata-X PRO cartridge (Phenomenex), the final product was eluted with 150 μL ethanol and reformulated with PBS/BSA (1.0% bovine serum albumin) solution for injection (11 KBq [^225^Ac]-PSMA-617 in 100 μL per mouse).

### Animal Studies

All animal studies were performed under the Guide for the Care and Use of Laboratory Animals through the Washington University Animal Studies Committee. The tumor model used for these studies were 22Rv1-Luc cells (a gift of Dr. Ken Pienta of Johns Hopkins University) grown in conditions specified by the American Type Culture Collection. Four million cells were implanted in a 1:1 mixture of cells in DPBS (ThermoFisher) and matrigel (Corning) into NCI Athymic NCR-nu/nu male mice (8wk; Charles River/NCI). Tumors were monitored by bioluminescence and caliper measurements, and used for treatment and imaging upon reaching approximately 500 mm^3^. A larger tumor volume than for traditional tumor control studies was chosen for the present investigations to preclude full local tumor control in the therapy group; reducing the variable of tumor size between groups.

Tumor bearing mice were anesthetized with isoflurane (2% mixture in 2L/min) and radioligand were delivered by injection into the retro-orbital sinus^24^. 300 nCi of [^225^Ac]-PSMA-617 was administered to each of the animals in the treatment group. For dynamic imaging, an animal from each treatment and control group were simultaneously injected with approximately 200 µCi on the bed of the dedicated small animal tomograph (microPET R4, Concorde Microsystems), in triplicate, and imaged for 30 minutes. All static images were acquired for 10 minutes at between 110-130 minutes post-administration. Acquisitions were recorded using an energy window of 350–700 keV and coincidence-timing window of 6 ns. The 30 minute duration datasets were histogrammed to 5 minute frames and were reconstructed using a *maximum a priori* MAP and 3D filtered back projection using a ramp filter with a cut-off frequency equal to the Nyquist frequency into a 128 × 128 × 63 matrix. Region of interest analysis was conducted in ASIPro (v6.3.3.0, Concorde Microsystems) and statistical analyses were performed in Prism (v8.0, GraphPad Software).

### Autoradiography and Immunohistochemistry

Fresh frozen sections of [^225^Ac]-PSMA-617 treated tumors were prepared and exposed, as previously ^25,26^. Briefly, 8 µm thick sections in O.C.T. Compound (TissuePlus, Fisher Healthcare) were fixed opposite a storage phosphor screen (MS, PerkinElmer) at -20° C for 3 d. Imaging and analysis were performed on the CyclonePlus and Optiquant (v4.1, PerkinElmer). Contrast limited enhanced histogram equalization was performed on the lossless image and displayed with an optimized colormap (Viridis^27^) using FIJI^28^.

Sectioned tissue slides were fixed briefly in 4% paraformaldehyde (Affymetrix), dried and stored. Slides were rehydrated before steaming in EDTA pH 8 (Invitrogen) for 40 min. Endogenous peroxidases were quenched with BLOXALL (Vector Labs), and the slides were blocked for 1 h with Serum Free Protein Block (Dako). Slides were incubated with antibodies directed against PARP-1 (Abcam ab194586). Staining was visualized with ImmPRESS Polymer detection kit and ImmPACT DAB (Vector Labs).

## RESULTS

### Radiochemistry

[^18^F]-PARPZ was synthesized from a 4-fluorobenzaldehyde precursor using an improved procedure for greater yeild^21^. We achieved a radiochemical yield of 52% and radiochemical purity of 99.9%. at the end of synthesis. Specific activity of the tracer at preparation was 76 GBq/µmol (≥2000 mCi/µmol per synthesis). Baseline PARP-1 imaging was performed at 2 h post administration of [^18^F]-PARPZ one day following the delivery [^225^Ac]-PSMA-617 was administered 2 h prior to baseline PARP-1 imaging. We chose the 6 day treatment date for followup imaging as we would not expect significant differences between saline control and [^225^Ac]-PSMA-617 treated 22Rv1 tumors at this interim time. The half-life of Actinium-225 is 10 days, and the energy deposited from the initially localized radiotherapy at day 6 will be roughly 35% of the total dose. Indeed tumor volumes between the groups were not significantly different at either imaging dates.

### [^18^F]-PARPZ PET

We performed PET imaging of treated and control groups at day 1 and day 6 post-administration. Figure 2 shows coronal slices (left) and maximum intensity projection (right) images for representative control untreated (top) and [^225^Ac]-PSMA-617 treated (bottom) animals from the two imaging time points. As expected, hepatobiliary clearance resulted in a majority of PARP tracer activity in the intestine at the imaging timepoints indicated in both baseline and treatment PET acquisitions. Distinction of the tumor can be made in all subjects using [^18^F]-PARPZ (as annotated with an arrow, Figure 2A-D). Qualitatively, there are no notable differences in tracer distribution or tumor delineation between the control and treated subjects at day 1 or the untreated animals at day 6. Prominent uptake in the treated tumor in the latter (day 6) imaging session (Figure 2D) is observed, and more clearly resolved in the projection imaging data (Figure 2H).

**Figure 2:**
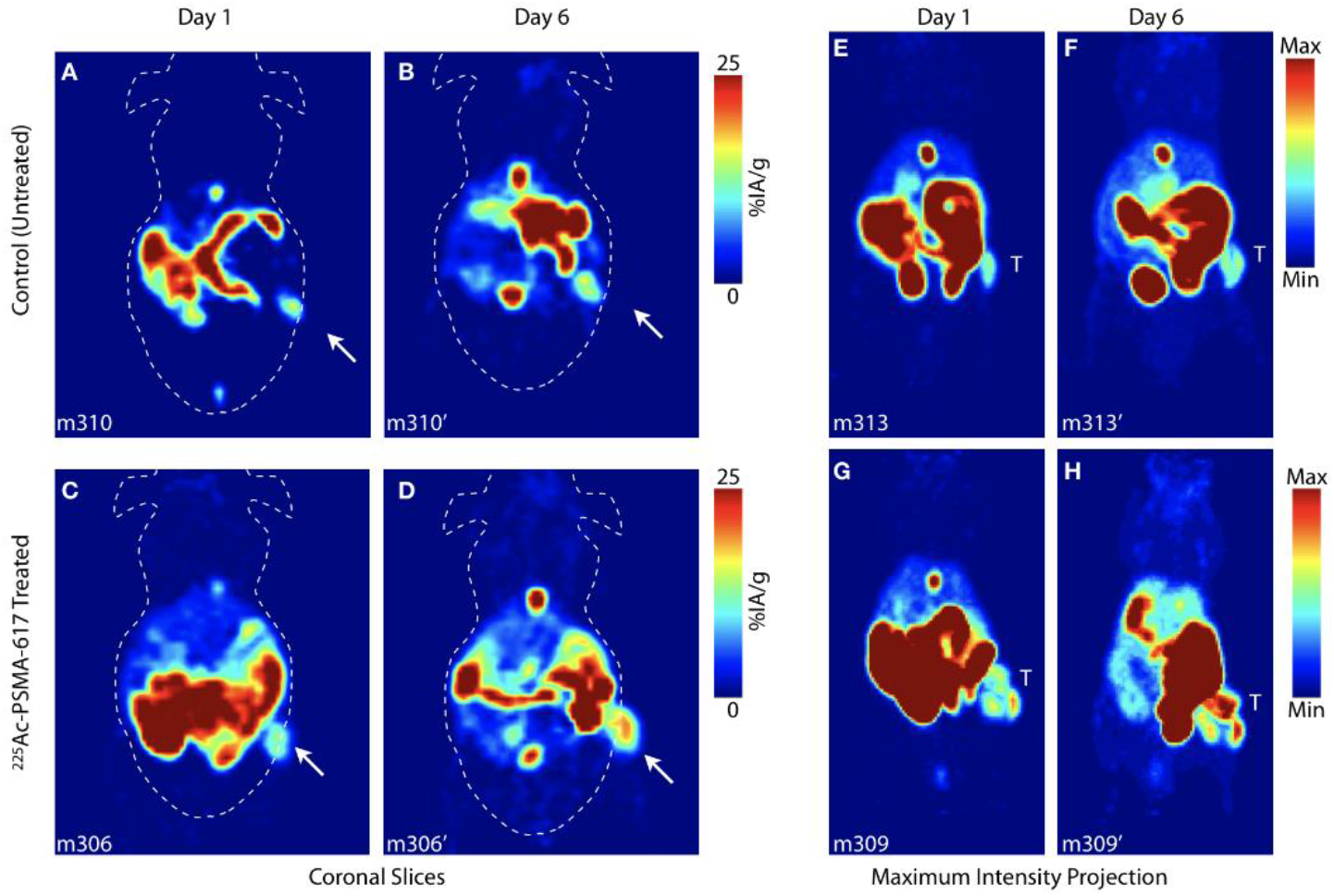
[^18^F]-PARPZ PET Imaging of Control and Alpha Particle Irradiated Tumors. Coronal slice of representative control mouse bearing 22Rv1-Luc xenografts at A) Day 1 and B) Day 6. Whole-body coronal PET slice of [^225^Ac]-PSMA-617 treated subject at C) Day 1 and D) Day 6. Distribution to the intestine (and gall bladder) recapitulates our previous work21; and tumor is delineated as indicated (arrow). Representative Day 1 and Day 6 whole body maximum intensity projection (MIP) data for control (top) and treated (bottom) groups; tumor denoted (T).

Quantitation of [^18^F]-PAPRZ PET uptake in tumors was assessed noninvasively and compared across control and treatment groups between the two imaging timepoints (Figures 3A and 3C). No significant difference in the tracer localization was noted between control and [^225^Ac]PSMA-617 groups at the initial baseline (1 day post-administration) scan. Likewise, no difference in mean or maximum tumor uptake was noted in the untreated group between day 1 and day 6 timepoints. In contrast, there was both a significant difference in PARP-1 tracer localization (mean percent injected activity per gram) for the treated group between the initial and latter scan dates (P<0.01); as well as a measurable difference between the control and treated tumor uptake at day 6 (P<0.05). Maximum tumor uptake as percent injected activity per gram was also significant between the treated animals across the two imaging timepoints (P=0.05).

**Figure 3:**
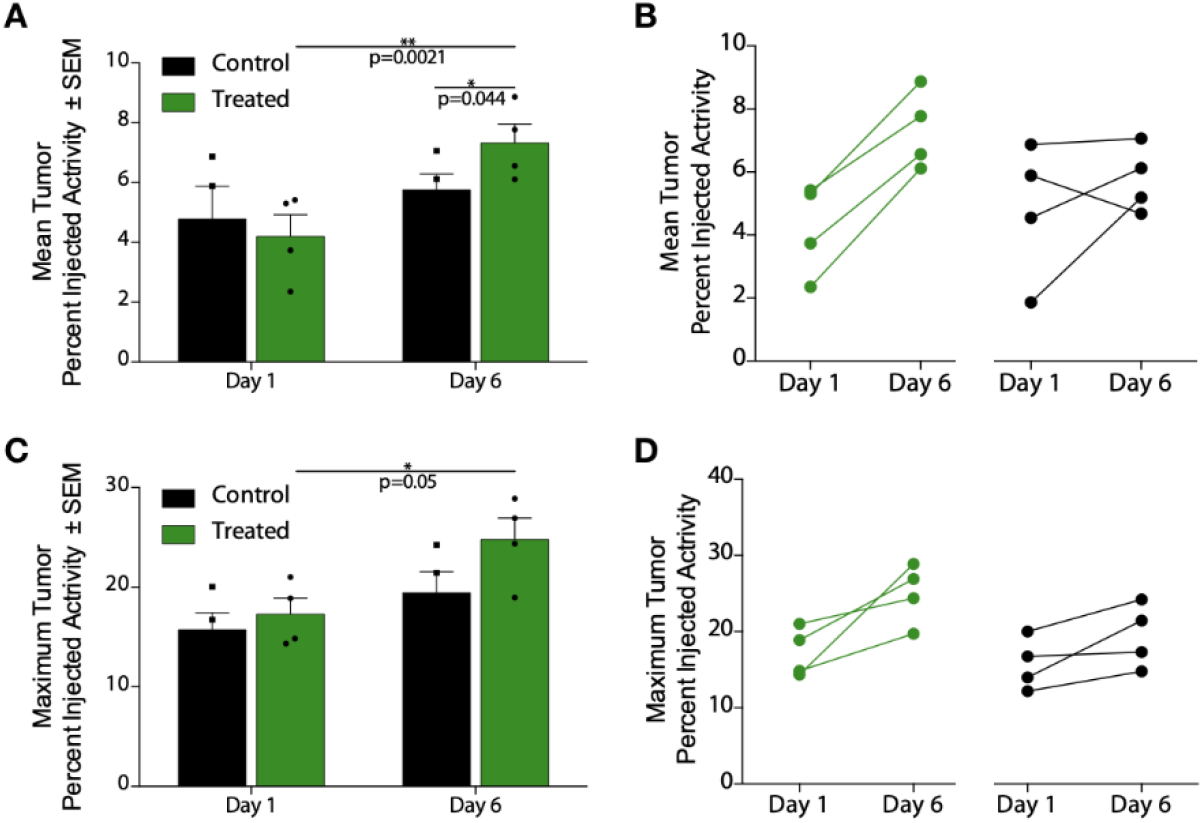
[^18^F]-PARPZ Uptake and Response Data: Noninvasive PET imaging analysis was used to measure [^18^F]-PARPZ uptake in treated and control groups. A) Mean percent injected activity per gram (%IA/g) in the tumors did not significantly vary at the early imaging time point. After 6 days of [^225^Ac]-PSMA-617 decay, the mean %IA/g significantly increased in the treated group (P<0.05). The two groups can be distinguished at this later time point by the mean uptake values (P<0.005). B) Before-after plot of the individual changes in replicates’ mean uptake values. C) Maximum tumor voxel %IA/g is plotted, showing an increase for the treated animals between Day 1 and Day 3 (P<0.05). A trend for increased maximum [^18^F]-PARPZ is present between the control and treated groups at Day 6, but is not significant. D) Changes in individual replicates’ maximum %IA/g.

The change in mean and maximum [^18^F]-PAPRZ uptake on an individual subject basis is shown in Figures 3B and 3D. The maximum intensity values increase in nearly all tumors investigated. We surmise that the increase in this metric is a function of both therapy induced DNA damage induced expression of PARP-1, as well as increased tumor cell PARP-1 expression in controls resulting from tumor progression.

A subset of animals were imaged on-camera upon tracer administration at the 6 day post-treatment imaging session in order to investigate the kinetics of tumor localization. The (mean) percent injected activity per gram accumulation in the tumor was recorded across 180 second frames out to 30 minutes. Two representative animals from each saline control and treated groups were imaged together on the imaging bed, and are plotted in Figure 4. These kinetic scans show more rapid uptake at an increased magnitude for the [^225^Ac]-PSMA-617 treated subjects.

**Figure 4:**
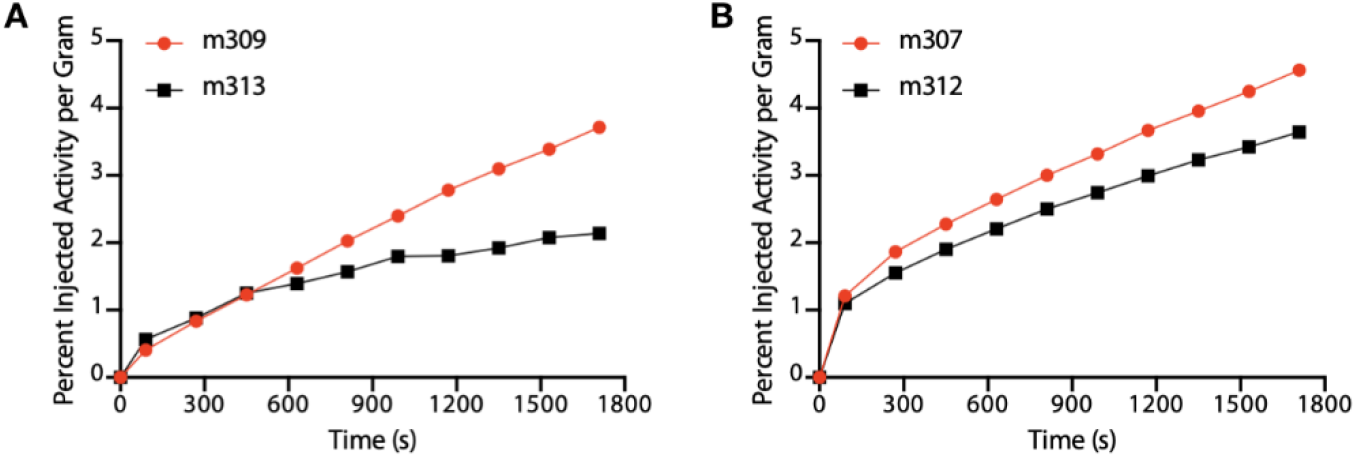
Rapid Uptake in [^225^Ac]-PSMA-617 Treated Tumors: The tumor specific activity concentration of the PARP-1 tracer was determined in treated and control mice co-injected on camera. The results indicate a trend of more rapid uptake of [^18^F]-PAPRZ in tumors treated with alpha particle emitting [^225^Ac]-PSMA-617 (red) over control saline treated subjects (black).

### Autoradiography and Immunohistochemistry

The dissected tumor tissue were sectioned and evaluated for [^225^Ac]-PSMA-617 signal by autoradiography in the treatment group and both groups were immunostained for PARP-1 (Figure 5). These tumors expressed PARP-1 either throughout, or in defined, non-stromal areas. The areas for which PARP-1 staining was present generally correlated with [^225^Ac]-PSMA-617 localization by phosphor autoradiography. No significant difference in staining intensity was observed between the control and treated groups. In the control group there was a trend towards more mild PARP-1 staining.

**Figure 5:**
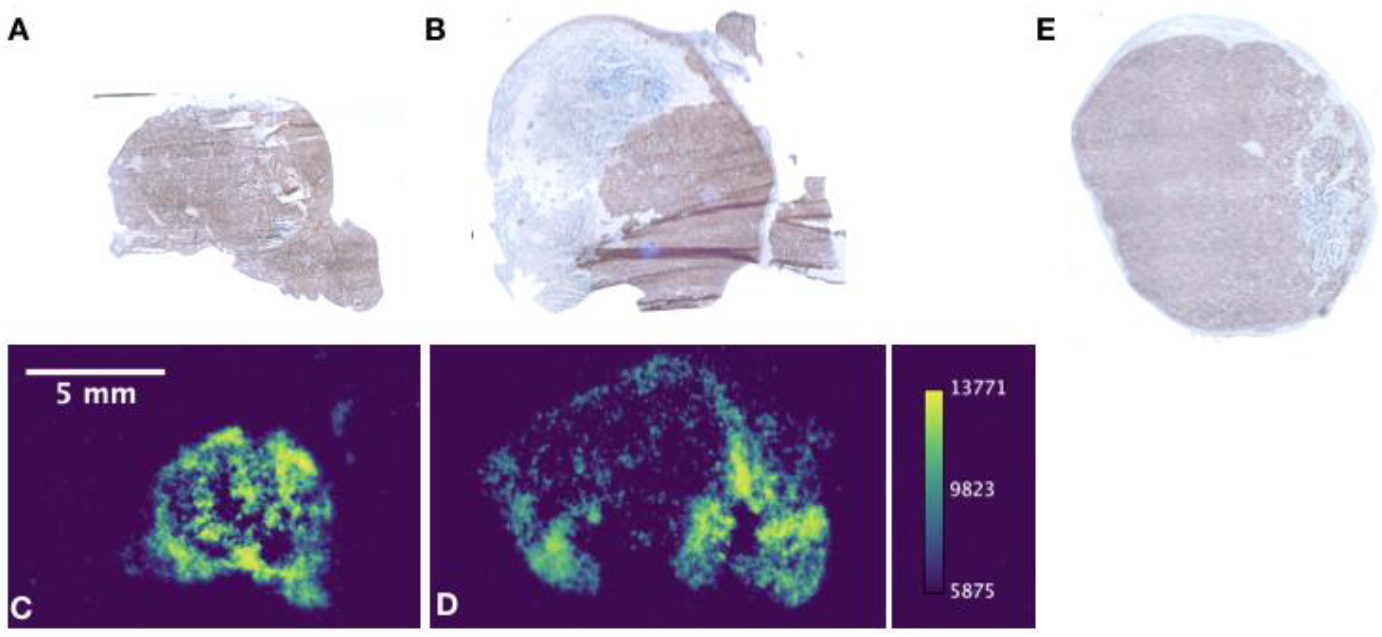
Immunohistochemistry and Autoradiography: Staining for PARP-1 was performed in tumors excised from treated and control animals, while autoradiographic images of tumor sections were acquired for [^225^Ac]-PSMA-617 in the treatment group. The areas of intense PARP-1 staining in treated tumors (A, C) and alpha particle emitting therapeutic (B, D) generally correlate. Staining of tumors in the control saline-treated group (E) tended to be less intense, but there was no significantly discernible difference in PARP-1 expression by IHC at this late sacrifice time point.

## DISCUSSION

We have evaluated the utility of a PARP-1 tracer to delineate responses to targeted alpha particle radiotherapy in a mouse model of prostate cancer. This work shows that we are able to quantitatively distinguish [^18^F]-PARPZ uptake in tumors treated with an alpha particle emitting targeted radiotherapy. The general focus of extant PARP-1 positron emitting radiotracer work has been to evaluate and validate PARP-1 expression for eventual clinical use^19,20,29^. This may provide the capacity to characterize PARP-1 expression in patients which may help to guide selection and dosing of PARP inhibitors, of which several are clinically approved for various indications^30^. Previous investigations of positron emitting PARP-1 tracers to noninvasively measure the activation of DNA repair cascades invoked in tumors by external beam X-ray irradiation have been published using [^18^F]-FluorThanatrace and [^18^F]-Olaparib^11,29^. A direct comparison between the various PARP-1 imaging agents published to date is difficult, as various oncologic models and therapies have been utilized^11,21,31–34^. What distinguishes this work is the use of [^18^F]-PARPZ, with minimal degradation or metabolic products^21^, comparatively very high tumor localization (at roughly 10% Mean IA/g and 20% Maximum IA/g; Figures 2,3), sustained uptake (Figure 4) and the use of potent alpha particle radiotherapy directed to a widely investigated prostate cancer radioligand target. While it is difficult to draw significant conclusions regarding the differences in PARP-1 expression following treatment, we were able to generally correlated PARP-1 expression IHC with areas of radiotherapeutic localization by autoradiography.

The differences in tracer kinetics in the tumors between treated and control groups (Figure 4) is an interesting observation. Our results demonstrate greater and more rapid localization of [^18^F]-PARPZ in [^225^Ac]-PSMA-617. In further planned studies, we intend to determine if the input characteristics of PARP-1 imaging tracer is indicative of treatment response. Limitations of this study include the limitation to a high PARP-1 expressing prostate cancer cell model. A quantitative evaluation of the relative sensitivities of detection of PARP-1 expression across cell lines by [^18^F]-PARPZ is warranted to fully evaluate this promising new tool.

## CONCLUSIONS

We are able to measure significant increases in PARP imaging tracer localization to targeted *α*-therapy treated tumors in a model system of prostate cancer. The improved imaging characteristics and stability of [^18^F]-PARPZ coupled with our institution’s successful translation and implementation of a previous-generation agent, [^18^F]FTT, lay the ground-work for the initiation of clinical investigation of DNA damage response imaging in patients treated with both approved (Radium-223 dichloride) and investigational alpha particle radiotherapy.

## ACKNOWLEDGEMENTS

This work was supported by R01-CA201035, AND R01-CA229893. The Siteman Cancer Center is funded by NCI Cancer Center Support Grant #P30 CA091842.

